# DENDRO: Recovery and denoising of whole-tree dendritic voltage from 2D voltage movies

**DOI:** 10.64898/2025.12.01.691576

**Authors:** Benjamin Antin, Pojeong Park, Amol Pasarkar, Utku Ferah, Chase King, Adam Cohen, Liam Paninski

## Abstract

The ability to image voltage at high spatiotemporal resolution across an entire dendritic tree would represent a major advance in systems and circuit neuroscience. Recent advances in genetically encoded voltage indicators (GEVIs) have brought this possibility closer to reality. However, due to fundamental tradeoffs between imaging speed, resolution, SNR, and volume, this goal has remained out of reach. Here we develop a computational method that fuses 3D anatomical information with 2D voltage video data, yielding full time-varying 3D voltage estimates. Our method, termed DENDRO, comprises two steps. In step one, we use the anatomical data to build a microscope model which maps from voltages along the tree to observed fluorescence at the imaging plane. By exploiting local spatial smoothness of the voltage signal, we parameterize the voltage signal using a set of local basis functions, which reduces the dimensionality of the problem and allows us to approximately invert the microscope model. This step leverages spatial but not temporal smoothness of the underlying signal and yields noisy 3D estimates. In step two, we train a lightweight self-supervised neural network to perform spatiotemporal denoising of the inferred voltages. On simulated data, we find that DENDRO is able to recover voltages at high accuracy across an entire dendritic tree. On real voltage movies from hippocampal slices, DENDRO recovers known dendritic phenomena at single trial resolution and millisecond time-scales, and allows visualization of backpropagating action potentials in 3D.

## 1 Introduction

The ability to record membrane potential along entire dendritic trees would be a powerful tool for understanding dendritic computation. Recent developments in genetically-encoded voltage indicators (GEVIs) [Abdelfattah et al., 2023; Evans et al., 2023; Hao et al., 2024] and microscopy techniques [Kazemipour et al., 2019; Wu et al., 2020; Wong-Campos et al., 2023] have brought this long-held goal closer to reality. However, voltage imaging requires very fast frame rates: at least 1 KHz to capture the millisecond-scale dynamics of action potentials. Therefore, because of fundamental tradeoffs between imaging speed, resolution, SNR, and field of view, conventional microscopy approaches do not yet deliver volumetric imaging of an entire dendritic tree at the speed and SNR required to recover the full spatiotemporal dendritic voltage signal.

In this study, we introduce a novel computational framework, DENDRO (DENdritic Denoising and Recovery), which sidesteps this tradeoff by combining 2D voltage movies, such as those obtained by widefield one-photon (1p) imaging, with known dendritic anatomy, to yield estimates of time-varying 3D voltage signals across large sections of dendritic trees at millisecond resolution. Our method proceeds in two steps: first, we use the known anatomy of the tree to formulate the problem of voltage inference as a linear inverse problem. This step proceeds frame-by-frame, and exploits spatial but not temporal smoothness of the underlying signal. The second step exploits spatiotemporal smoothness by training a graph-structured neural network to denoise voltage estimates from the first step.

To validate DENDRO, we used morphological reconstructions of CA1 pyramidal neurons to generate realistic voltage imaging movies. We found that our model could recover ground-truth voltage signals, significantly improving estimates compared to using the voltage movies on their own. We then applied our framework to 1p voltage movies in acute slices from Park et al. [2025] and recovered 3D estimates of back-propgating action potentials (bAPs) and plateau potentials.

## 2 Related work

Previous attempts at computational recovery and denoising of dendritic voltage largely have focused on simulated data (rather than real data, as demonstrated here), and have either relied on sampling methods [Huys and Paninski, 2009; Omori and Hukushima, 2016; Sun et al., 2019], which scale poorly to large real dendritic voltage datasets, or have approximated dendritic voltage using a Kalman smoother model [Paninski, 2010], which oversmooths spikes. Pnevmatikakis et al. [2012] presents a related model for denoising calcium dendritic measurements; however, calcium signals are much slower and higher-SNR than voltage signals and therefore pose rather different computational and statistical challenges than those considered here.

Another line of related work centers on denoising voltage or calcium movies themselves, without considering dendritic anatomy. These methods all rely on some variant of self-supervised blind spot denoising, in which a neural network is trained to predict the value of a held-out center pixel given the surrounding pixels [Batson and Royer, 2019; Krull, Buchholz, and Jug, 2018]. Lecoq et al. [2021], X. Li, Zhang, et al. [2021], and X. Li, Y. Li, et al. [2023] designed networks to predict held-out frames of calcium imaging movies, but their approach did not leverage spatial structure within each frame. Eom et al. [2023] and B. Wang et al. [2024] augmented the blind-spot approach to include spatiotemporal context for each held-out pixel, leading to improved performance on the quickly-varying signals present in voltage imaging data. Y. Wang et al. [2025] implemented a spatiotemporal sampling strategy during training which avoided the need for 3D convolutions, resulting in very fast inference speed. However, these video denoising methods do not on their own allow the extraction of 3D signals from 2D videos. DENDRO extends these methods to solve the 2D-to-3D inference problem.

## 3 Methods

We make two main methodological contributions to the problem of dendritic voltage inference. We begin by posing the problem of voltage inference as a linear inverse problem. To do so, we use the known dendritic anatomy to compute a set of smooth 3D basis functions, which are transformed by the microscope point spread function (PSF) to yield observed fluorescence at the imaging plane. Second, we design a custom neural-network architecture to perform self-supervised denoising of voltage on dendritic trees. A high-level overview of our approach is shown in Figure 1. In all our experiments, we found it beneficial to apply penalized matrix decomposition [Buchanan et al., 2019; Pasarkar et al., 2023] to the raw functional video data, as a lightweight spatial denoiser and dimensionality reduction before voltage inference.

**Figure 1.**
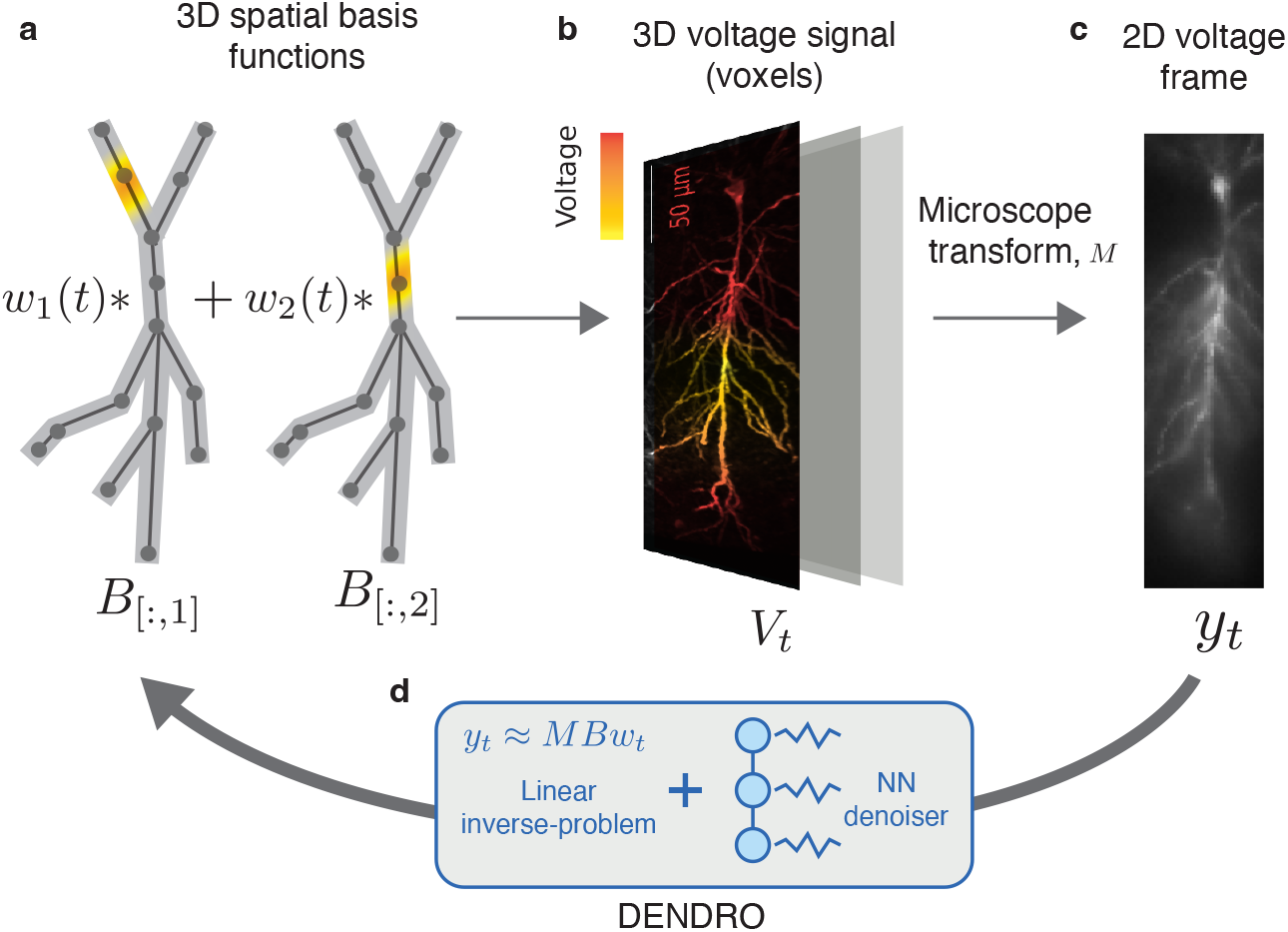
**a**, Coarse dendritic anatomy is represented by a graph (dark grey circles and lines) and fine-grained anatomy is represented by a 3D array of voxels (light gray shaded region). A smooth, locally supported 3D basis function is placed at each node of the graph. **b**, At a given frame *t*, the basis *B* is weighted by a vector of weights *w*_*t*_, forming a smooth 3D voltage signal *V*_*t*_ = *Bw*_*t*_ across the whole tree. **c**, This 3D voltage signal is passed through a microscope model *M* and corrupted by noise to generate a given frame *y*_*t*_. **d**, DENDRO recovers 3D voltage signals by approximately inverting the transformation from node weights to pixel fluorescence, and denoising those estimates with a neural network.

### 3.1 Formulating voltage inference as a linear inverse problem

To successfully tackle the voltage inference problem on real data, we need an effective low-dimensional parameterization of smooth 3D voltage signals. Multicompartment models, such as those used in simulation studies of dendritic voltage denoising [Paninski, 2010; Sun et al., 2019], make the assumption that voltage is spatially constant within a compartment. While this assumption works well for biophysical modeling, it leads to a very high-dimensional inverse problem (many compartments *×* many time points). To reduce this dimensionality, we compute a library of smooth 3D basis functions with local support, which can be used to approximate 3D voltage signals.

We assume that structural imaging gives us access to two types of data. First, an undirected graph *G*, whose nodes and edges capture the coarse morphology of the tree (Figure 1a, dark grey dots and lines). Second, a voxelized segmentation represented by a sparse 3D array, which captures more fine-grained aspects of the tree morphology, such as branch thickness (Figure 1a, light gray shaded region).

Letting *P* be the number of nodes in *G*, we place a smooth, locally supported basis function at each node (Figure 1a, bright orange regions). We use simple graph-triangular splines for these basis functions; see subsubsection A.4.1 for details. We additionally define a *P*-length vector *w*_*t*_, where a single element *w*_*j*_(*t*) defines the relative weighting of node *j* at timestep *t*. Letting *Q* denote the number of voxels in the segmented dendritic morphology, the basis functions form the columns of a *Q × P* matrix *B*, so that at time-step *t*, the voltage vector *V*_*t*_ = *Bw*_*t*_ is a smooth 3D signal composed of weighted basis functions (Figure 1b).

Having defined the mapping *B* from node weights to smooth 3D signals, we now need a forward mapping from 3D signals to 2D observed fluorescence. We define a second matrix *M*, which captures the PSF of the microscope. *M* is a *D × Q* matrix, where *D* is the number of pixels in the voltage movie and *Q* is the number of voxels in the neuronal morphology. The matrix *M* performs the (linear) blur-and project operation, in which each plane of the anatomy stack is convolved with a Gaussian kernel. The width of this kernel increases when moving away from the focal plane. See subsubsection A.4.2 for details.

Combining the microscope matrix *M* and the basis matrix *B*, we can model the voltage movie as

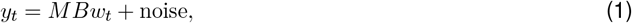

where *y*_*t*_ is the *D*-vector (image) of fluorescence values recorded at each timestep. However, this model does not yet capture an important aspect of the experiment – namely that we do not expect indicator expression to be perfectly uniform throughout the cell, nor do we expect observed fluorescence to be zero when the voltage is at baseline. To address this issue, we add a gain term to the model for each voxel. To capture fluorescence when the cell is at rest, we further add an offset to each pixel. With these terms added, the model becomes:

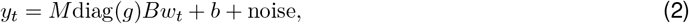

where *g* is a *Q*-vector representing the gain for each voxel, and *b* is a *D*-vector representing the offset for each pixel.

### 3.2 Solving the linear inverse problem at each frame

For a movie indexed by time-steps *t* = 1, …, *T* the model above can be fit by alternating minimization over the node weights *w*_*t*_, the gains *g*, and the offsets *b*:

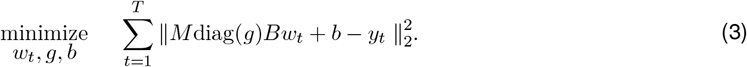

In practice, we found that solving for *w*_*t*_ was ill-posed for the microscope matrices *M* we consider in our experiments; because *M* encodes a 3D to 2D transformation, it is ill-conditioned. We therefore added a graph-Laplacian regularizer to (modestly) encourage spatial smoothness within each frame:

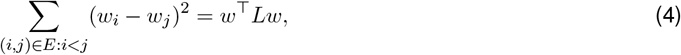

where *E* denotes the edge-set of 𝒢 and *L* the graph Laplacian.

This leads to the following regularized objective to infer node weights *w*_*t*_ for each timestep: *w*_*t*_ minimize

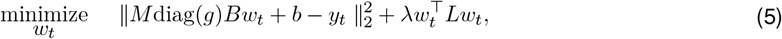

which has a simple closed form solution when *g, b* and *λ* are held fixed. In order to choose an appropriate value for *λ*, we use generalized cross validation (GCV) [Golub, Heath, and Wahba, 1979], which forms a computationally efficient estimate of leave-one-out cross validation for underconstrained problems. With this regularizer in place, we can alternately optimize over *w*_*t*_, *g, b* to fit the model.

Since fitting *w*_*t*_, *g* and *b* jointly is non-convex, this model is sensitive to initialization. We found that initializing *g* and *b* with the voltages *V*_*t*_ held spatially constant created more stable solutions. Following this initialization step, we do alternating minimization on *w*_*t*_, *g, b*, where each update to *w*_*t*_ uses GCV to find the optimal penalty parameter *λ*. Finally, we observed that the penalty parameter *λ* selected by GCV tended to over-smooth spikes, since traveling wave-fronts are less spatially smooth than subthreshold events. To ameliorate this effect, we simply divided the GCV estimated penalty with a large constant factor at the last iteration. While this increased the noise present in the estimated weights, our neural network denoiser (discussed in the next section) could easily compensate for the increased noise level. The full algorithm for fitting *w*_*t*_, *g*, and *b* is shown in subsubsection A.4.3.

### 3.3 Designing a graph-structured denoising network

The steps described in subsection 3.2 yield time-varying node weights *w*_*t*_. Since we have so far used spatial, but not temporal information, the estimates at each frame suffer from temporal noise. We therefore sought to design a spatio-temporal denoiser to further improve the SNR of our voltage estimates. As detailed in prior work, denoising voltage on a dendritic tree is a difficult statistical problem, because there are multiple smoothness levels present in the data [Sun et al., 2019]: in the sub-threshold regime, voltage-gated channels are inactive, and therefore the dynamics are quasilinear and relatively smooth. These slowly-varying voltage dynamics can be effectively handled by a Kalman smoother model [Paninski, 2010]. However, when the cell enters the spiking regime, voltage dynamics become highly non-linear and evolve much faster. Therefore nonlinear denoising methods are required; here we use a neural network denoising approach.

Taking inspiration from blind-spot denoisers for image and movie data, we designed a self-supervised denoising network to operate on graph-based time series data. Our main insight is that the masked convolutions used in previous blind-spot approaches [Eom et al., 2023; B. Wang et al., 2024] can be generalized to operate on graphs. Figure 2d shows the two fundamental operations of our architecture: a spatial blind-spot operation implements message-passing on the graph, but disallows a node from passing messages to itself. A temporal blind-spot operation aggregates information from adjacent timepoints. We combined these two operations into a “graph-temporal blind-spot layer” (Figure 2e) and then stacked these layers following the architecture suggested by Honzátko et al. [2020]. The full architecture is described in subsection A.5.

**Figure 2.**
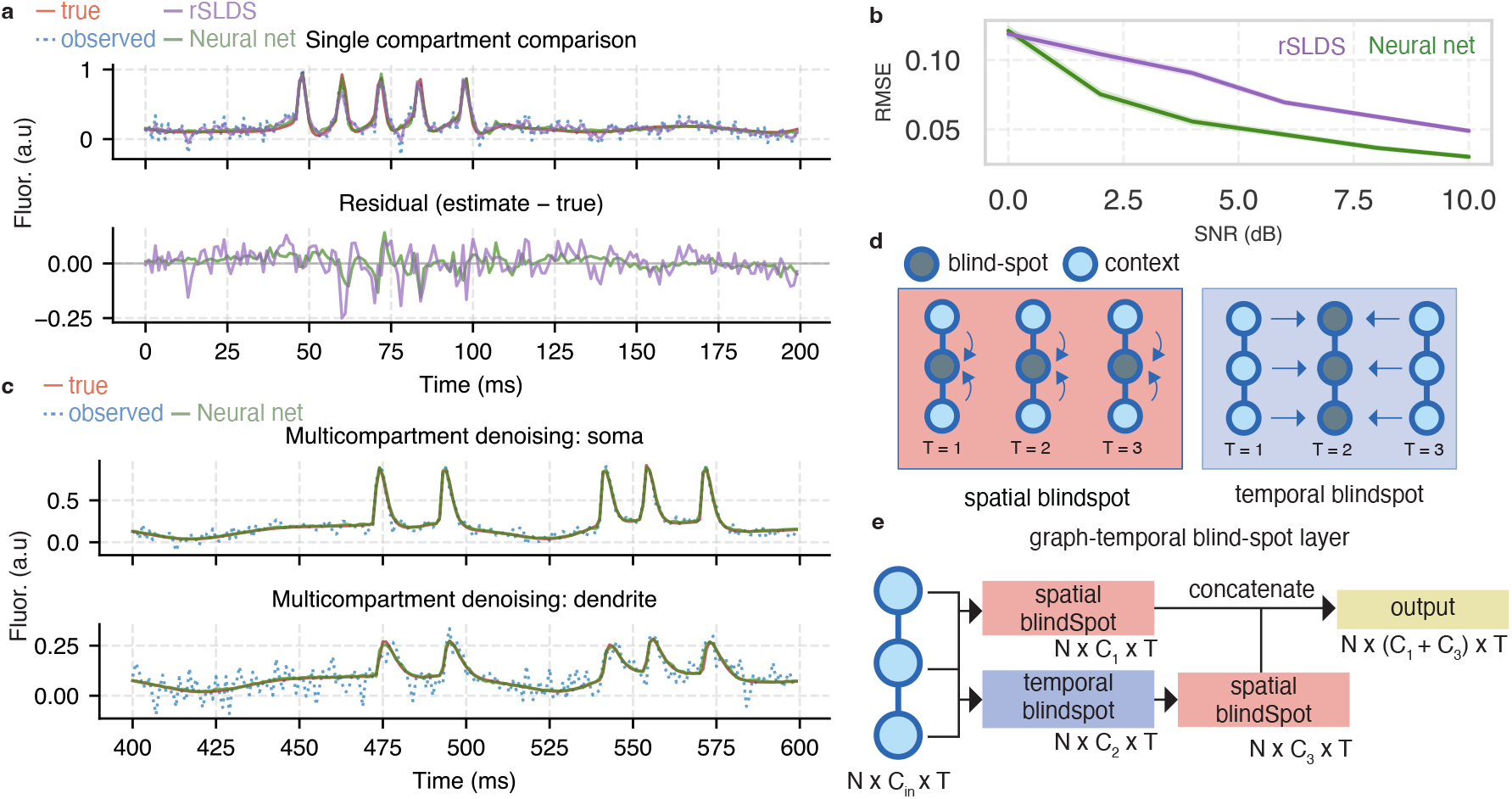
Blind-spot denoisers are effective for denoising dendritic voltage. **a**, Comparison of blind-spot model to rSLDS on single-compartment simulated data. Top: ground truth, observed, rSLDS denoised, and blind-spot denoised traces. “Neural net” here refers to a temporal-only version of the blind-spot architecture (since this is a single-compartment simulation). Bottom: Residuals (estimates -true) for rSLDS and temporal blind-spot network. **b**, Comparison of rSLDS and blind-spot model (NN) on single-compartment data across various noise levels. Shaded area is +/− 95% CI bootstrapped over 10k timesteps (difficult to see due to overlap). **c**, Demonstration of the graph-based blind-spot model denoising data from a multicompartment simulation. “Neural net” here refers to the full graph-based spatiotemporal model. Top: soma compartment. Bottom: distal dendritic compartment. The SNR is much lower in the dendritic compartment, but the network effectively pools information across space to achieve good performance. **d**, The two fundamental operations in our architecture. A “spatial blind-spot” operation passes messages between nodes, but disallows a node from passing messages to itself. A temporal blindspot block uses a masked convolution to pool information from neighboring timepoints. **e** These two operations are combined in a “graph-temporal blind-spot” layer, which forms the core of our architecture; see subsection A.5 for details.

Finally, to capture fast transients due to spikes, we re-combined the blind-spot denoiser output with the raw observed data, using the adaptive Gaussian combination rule described by Laine et al. [2019]. As desired, we observed that the model learned to upweight the input data during spikes (difficult to predict events) and downweight the input data during subthreshold periods (easy to predict events) (Figure S1c,e). See subsection A.5 for details.

## 4 Results

### 4.1 Blind-spot denoisers are effective for denoising dendritic voltage

We began by verifying that a temporal-only blind-spot model could effectively denoise single-compartment data. We used the Python package Jaxley [Deistler et al., 2024] to simulate a Hodgkin-Huxley model driven by a randomly varying current which elicited both sub-threshold dynamics and spikes. To accurately capture the transformation between voltage and fluorescence, we built a four state GEVI model, with trainable parameters that controlled the transitions between states. We used a dataset containing paired voltage clamp and fluorescence recordings to train the parameters of the GEVI model, by minimizing the mean-squared error between the predicted fluorescence and the observed fluorescence, given the patch data as a ground-truth voltage measurement (Figure S2a). After training, we found that the trained model was able to faithfully reproduce the recorded fluorescence traces during spikes elicited by current steps (Figure S2b). We fixed the GEVI parameters to their trained values and kept these values fixed for all simulated validation experiments – see subsubsection A.6.1 for details. We used 10k time-steps for all simulations, since our real datasets are 10k frames in length.

On single-compartment simulations, we benchmarked our temporal blind-spot approach against a recurrent switching linear dynamical system (rSLDS)^1^, using the same hyperparameters as used for single-compartment experiments in Sun et al. [2019]. We found that the blind-spot model outperformed the rSLDS uniformly across the physiologically relevant SNR range (Figure 2a,b).

Having established the viability of the blind-spot approach, we applied our graph-based denoiser to data from a multicompartment model. Since our real data experiments are on voltage movies of CA1 pyramidal neurons, we tuned our simulations to match these data using the multicompartment model described in Park et al. [2025]. After simulating a current injection to the apical dendrite, our model recapitulated backpropagating action potentials (bAPs) which sometimes resulted in dendritic sodium spikes (dspikes). Training our graph-based architecture on data from 133 compartments across 10k timesteps took 4.5 minutes on a single consumer-grade GPU.

Figure 2e shows an example trace from the soma and from the distal dendrite in our simulations. In the traces shown, the bAPs are attenuated by the dendrite and do not result in dendritic spikes. In this instance, voltage denoising is especially difficult in the dendritic compartment, because the SNR is much lower than at the soma (the noise level is constant, but the spike is attenuated). We found that our graph-based denoiser was able to closely match the ground truth even in this challenging case by pooling information across space and time.

### 4.2 DENDRO recovers voltages with high accuracy on simulated dendritic voltage videos

Our experiments in the prior section validated the denoising step of our pipeline, but they did not include effects of a microscope model. To validate DENDRO’s ability to recover 3D voltages, we generated voltages using the multicompartment model described above and then built a microscope matrix *M* matched to the PSF parameters of the real data (see subsubsection A.4.2). By passing the voltages from the multicompartment model first through the GEVI model, and then through the basis matrix *B* and the microscope matrix *M*, we generated simulated 2D voltage videos of spikes propagating along a dendritic tree and added multiplicative gaussian noise with SNR matched to the real data (see subsection A.6 for details). An example simulated video can be found in the supplemental data^2^. To mimic the pipeline used on real data, we applied PMD as a preprocessing step to all of our videos before applying DENDRO.

Figure 3a shows the anatomy stack used in our simulation, with a set of nodes from a selected branch overlayed. The selected branch includes both in-focus and defocused regions. Figure 3b compares the mean image of the real voltage movie for the same cell, with the mean image of our simulated movie, showing a good qualitative match.

**Figure 3.**
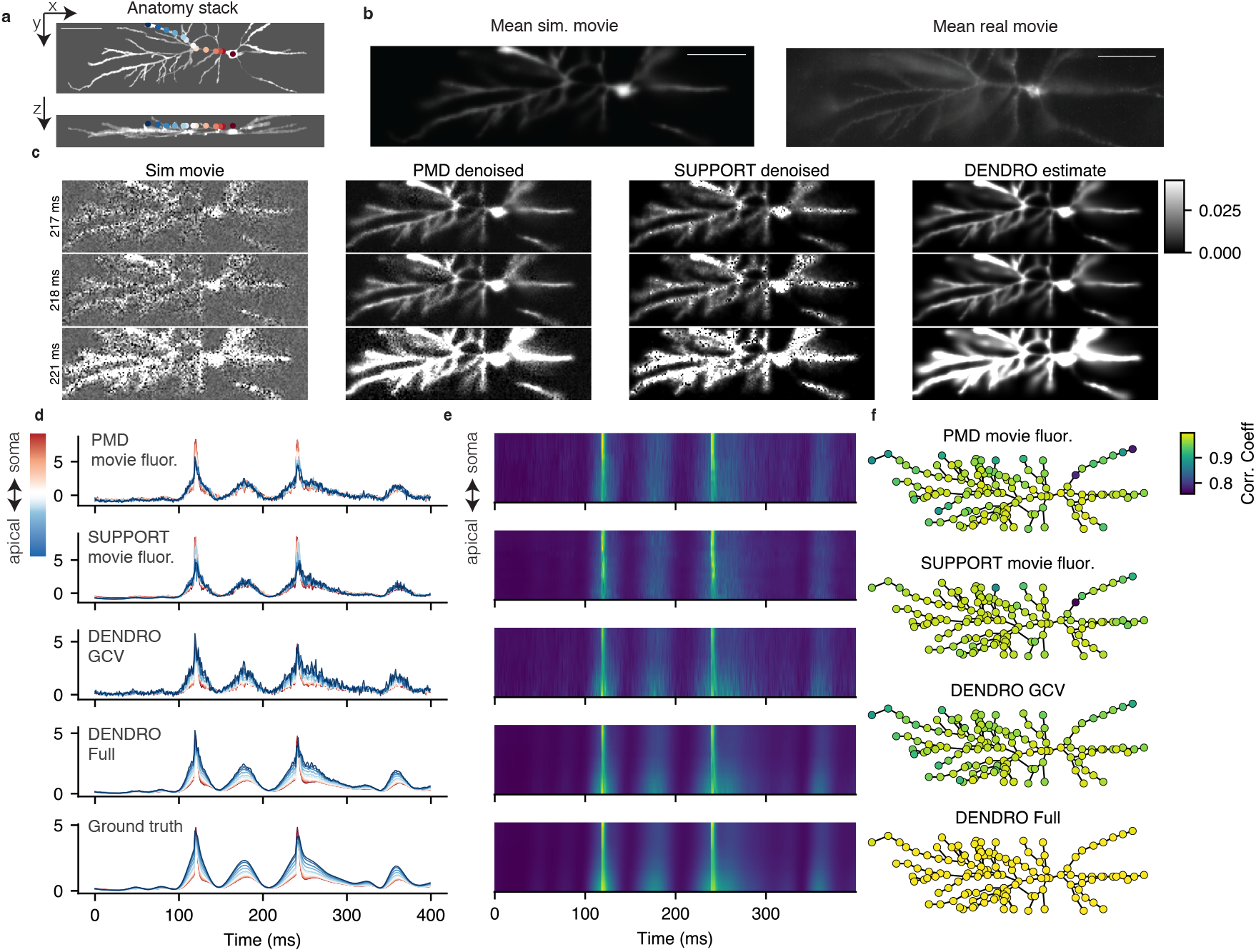
DENDRO recovers voltages with high accuracy on simulated dendritic voltage videos. **a**,XY and XZ projections of anatomy stack used to generate simulated voltage videos. Selected nodes are shown as overlayed dots. Current was injected at the most distal (dark blue) node. Scalebar is 75 *µ*m. **b**, Mean image from simulated movie and real movie. A noiseless simulated frame is generated as *y*_*t*_ = *MBw*_*t*_, where the weights *w*_*t*_ come from the biophysical model. **c**, Frames from raw and denoised movies during a spike. Rows show different time steps. From left to right: raw simulated movie, PMD denoised, SUPPORT denoised, and DENDRO estimate. The DENDRO estimate is formed as *ŷ*_*t*_ = *MBw*_*t*_ where *w*_*t*_ are the inferred node weights. **d**, Inferred traces for the nodes shown in **a** (trace colors match colors used for each node). “Movie fluor.” (top two panels) is calculated by computing Δ*F/F* for patches surrounding each node. From top to bottom: movie fluor of PMD denoised movie, movie fluor of SUPPORT denoised movie, DENDRO estimate with linear-inverse step only, Full DENDRO pipeline, ground truth. **e**, Same as **d** but visualized as heatmaps. **f**, Correlation coefficients visualized across the tree for the four different methods. When naively using movie fluorescence (top two panels) the correlation is lowest at branch end-points and thinner dendrites. DENDRO full (bottom panel) achieves high correlation regardless of spatial location.

We compared the ability of different approaches to estimate the voltage signal at each of the nodes shown in Figure 3a: (i) a naive “movie fluorescence” baseline that extracts Δ*F/F* traces from small patches around each node in the PMD movie, (ii) the same patch-based baseline applied to movies denoised by the SUPPORT network [Eom et al., 2023], (iii) the linear-inverse step of DENDRO alone (“DENDRO GCV”), and (iv) the full DENDRO pipeline with graph-based denoising. The entire DENDRO pipeline, including PMD and training the graph-based denoiser, took around 20 minutes per dataset (see subsection A.2); training SUPPORT took approx. 20 hrs.

Figure 3c shows frames from the simulated movie, along with denoised estimates from PMD, SUPPORT, and DENDRO. To form the DENDRO-estimated frames, we computed *ŷ*_*t*_ = *MBw*_*t*_ where *w*_*t*_ are the estimated node weights after denoising. As expected, we found that all three methods noticeably reduced noise of the movie – though DENDRO performed better because it leverages knowledge of the underlying anatomy.

For each node, we computed Δ*F/F* traces by averaging pixel values in a patch surrounding the node, and compared these to the DENDRO GCV and full DENDRO estimates as well as the ground-truth values (Figure 3d,e,f). DENDRO provided estimates which had noticeably reduced noise and improved match to the ground truth data.

This improvement is reflected in the correlation maps over the entire tree (Figure 3f): correlations between ground-truth and naive movie-fluorescence traces degrade for thin or defocused regions of the tree, while full DENDRO maintains uniformly high correlation throughout the morphology (PMD corr. = 0.96 +/− 0.03, SUPPORT = 0.97 +/− 0.03, DENDRO GCV = 0.97 +/− 0.02, DENDRO Full = 0.99 +/− 0.01).

### 4.3 DENDRO recovers realistic voltage motifs on real data

Having established DENDRO’s performance on simulated voltage movies, we applied it to two acute slice datasets. The data are described in detail in Park et al. [2025]. In brief, each recording includes a 1p widefield voltage movie, along with two-photon 3D anatomy reconstruction. For each 1p movie, we applied PMD as a preprocessing step (as we did with the simulated data), and then applied DENDRO.

Using the aligned two-photon anatomy stacks, we built basis matrices and microscope matrices for each cell. After running the linear-inverse step of DENDRO (in which we simultaneously estimate node weights, offsets, and per-voxel gains) we observed a close match between our learned microscope model and the average of the 1p voltage movies (Figure 4a,d middle panel).

**Figure 4.**
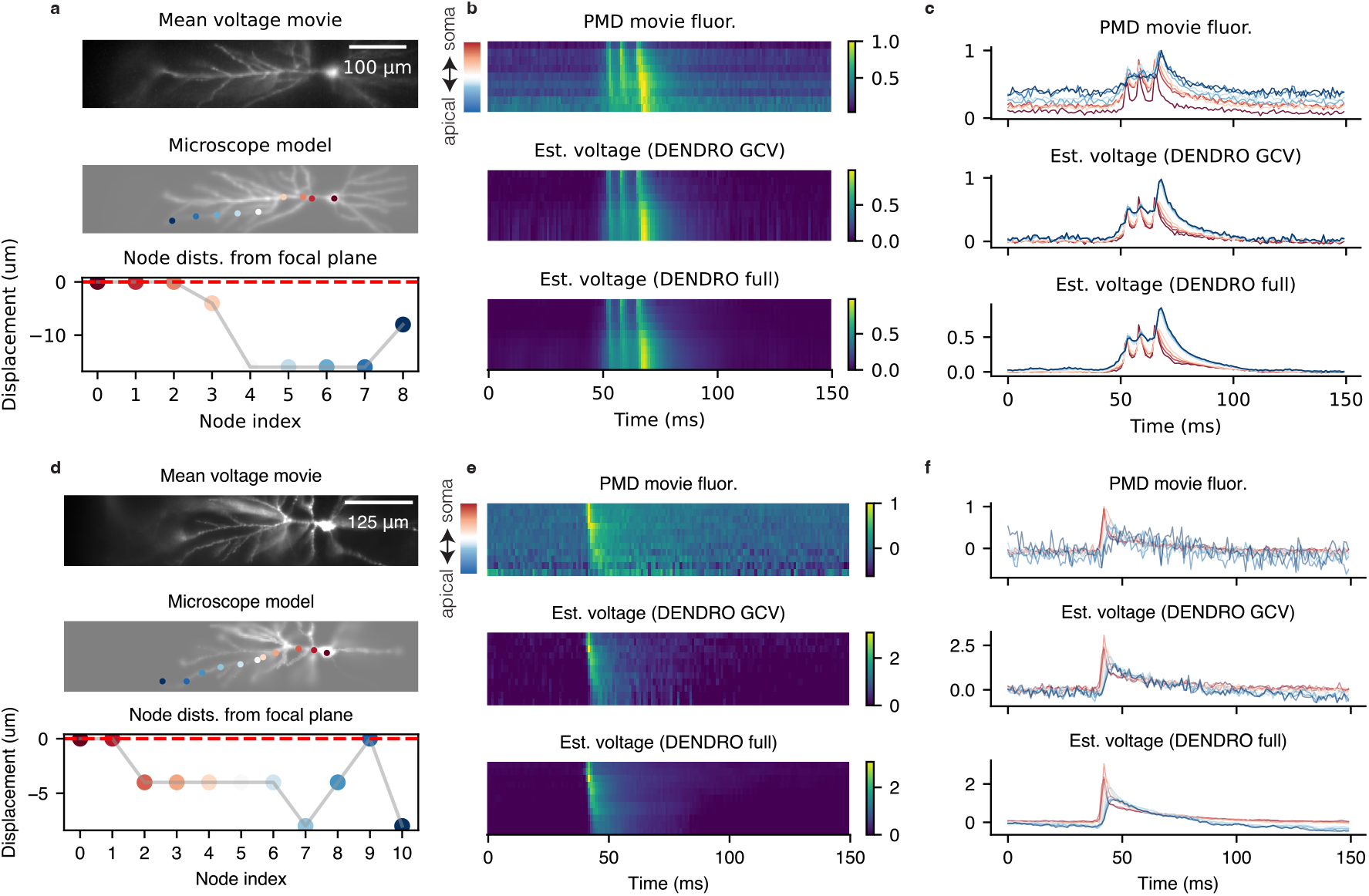
DENDRO recovers realistic voltage motifs on real data. **a**, Top: Mean of voltage movie. Middle: Example frame from microscope model, which captures blurring of defocused branches. Overlaid circles show selected nodes for a defocused branch. Bottom: displacements from the focal plane (dashed red line) for the overlayed node locations. **b**, Estimated voltage heatmaps for three different methods, for the nodes shown in **a.** Top: estimated by taking 3x3 patches from the PMD movie surrounding each node. Middle: “DENDRO GCV” refers to linear step of DENDRO without neural network denoising. Bottom: “DENDRO full” includes the neural net denoiser. **c**, Same as **b**, but visualized as traces. **d,e,f**, Same as **a,b,c** but for a second cell. See subsection A.8 for corresponding videos.

In the first dataset, as described in [Park et al., 2025], simultaneous optogenetic stimulation of the soma and the distal dendrites resulted in a motif of three backpropagating action potentials, the last of which was accompanied by a dspike. Applying DENDRO to this dataset recovered denoised estimates of this motif, even for branches which were not clearly in focus (Figure 4b, c, Supplemental Videos 4 and 5).

In the second dataset, brief stimulation of the soma elicited backpropagating action potentials without dspikes. Here too, DENDRO recovered estimates of the bAP wavefront which were qualitatively much lower noise than using the fluorescence from the PMD movie alone (Figure 4e,f, Supplemental Videos 4 and 5). In both cases, estimates obtained via DENDRO were significantly less noisy compared to using the PMD video on its own, which may be especially important when fitting biophysical models (see section 5).

## 5 Conclusion

We developed DENDRO, a two-stage pipeline for reconstructing and denoising 3D dendritic voltage signals from 2D voltage imaging data. Because the voltage signals of interest are typically much lower dimensional than the number of compartments in the tree, we introduce a low-dimensional parameterization, along with a simple spatial regularizer, which results in a tractable linear-inverse problem. Step 1 of DENDRO solves this regularized inverse problem separately for each frame of the movie. Given these 3D voltage estimates, DENDRO next performs further spatiotemporal denoising using a novel blind-spot graph neural network architecture.

Our approach to denoising represents a departure from previous blind-spot denoising methods applied to neural data. First, following [Laine et al., 2019], we incorporate data from the blind spot in our final estimates, leading to higher-quality denoising. Second, our architecture implements a graph-based blind-spot and performs denoising in node-space rather than in pixel-space. The choice to directly denoise the nodes of the graph has two main advantages for dendritic voltage imaging. First, it results in a network which is significantly more lightweight than those which have been used to denoise raw voltage movies: training a graph-based network on a dataset of 10k timesteps takes under 5 minutes with DENDRO, compared to 24 hours or more when denoising raw video directly [Eom et al., 2023]. Second, because the DENDRO network operates directly on the dendritic graph, it is able to handle non-uniform spatial sampling. This raises the exciting possibility of extending DENDRO to handle data from random-access or scanning microscopy methods [Kazemipour et al., 2019; Villette et al., 2019; Wu et al., 2020; Sims et al., 2024], simply by plugging in a new microscope matrix *M* .

There are some potential limitations to our work as presented. First, we do not have ground-truth data to validate the experiments on real datasets. Though it is not feasible to record ground-truth data across an entire dendritic tree, future work might be validated by dual-plane imaging, in which voltage inferences made on data from one focal plane could be checked against simultanous recordings from a second focal plane. Second, we observed during our experiments with real data that the model is very sensitive to errors in the microscope model, particularly misalignment between the voltage movie and the anatomy stack. Such alignment errors could skew voltage estimates if left uncorrected; an interesting direction for future work would be to automatically adapt the PSF parameters to the data.

It is natural to wonder whether our voltage estimates might be improved by directly incorporating a biophysical model. We chose a modeling approach that is somewhat agnostic to biophysics for a few reasons. First, errors in the specification of the biophysics model would likely lead to biases in the estimated voltages; our approach is somewhat more nonparametric and thus likely less biased (perhaps at the cost of increased output noise variance). Second, using a flexible neural-network denoiser means that DENDRO can potentially be applied to calcium or glutamate recordings with minimal tweaks. Still, directly exploiting biophysics for dendritic voltage recovery is a worthwhile direction for future work, especially given recent improvements in differentiable multicompartment simulation [Deistler et al., 2024].

Advances in dendritic imaging, when combined with computational recovery techniques like DENDRO, have the potential to unlock new avenues for studying dendritic computation. Direct knowledge of the voltage throughout a dendritic tree would enable automated fitting of multicompartment models [Huys, Ahrens, and Paninski, 2006; Deistler et al., 2024], which may allow for the study of cell-to-cell biophysical variability. Looking forward, we anticipate that DENDRO could be extended to reconstruct spatiotemporally varying dendritic signals *in-vivo*, ultimately providing a more complete picture of information flow in neural circuits.

## Acknowledgements

The authors would like to thank Marcus Triplett and Darcy Peterka for useful discussions. This work was supported by Chan Zuckerberg Initiative Dynamic Imaging grant 2023-321177 (A.E.C. and L.P.), the Kavli Foundation and the Gatsby Charitable Foundation grant GAT3708 (L.P.).

## APPENDIX A Appendix / supplemental material

### A.1 Animal ethics statement

All animal procedures adhered to the National Institutes of Health Guide for the care and use of laboratory animals and were approved by the Institutional Animal Care and Use Committee (IACUC).

### A.2 Computational resources

All experiments were run on a workstation with 128gb RAM, with 6 cores (12 threads) running at 3.6GHz. For training the denoising networks and running PMD, we use a single NVIDIA GeForce GTX 1080 Ti.

### A.3 Notation

| symbol | definition | shape | typical value |
| --- | --- | --- | --- |
| $T$ | Number of timesteps in voltage movie | scalar | 10000 |
| $D$ | Number of pixels | scalar | $\mathcal{O}(10000)$ |
| $P$ | Number of nodes | scalar | 100-300 |
| $Q$ | Number of foreground voxels | scalar | 3000-10000 |
| $M$ | microscope matrix | $D \times Q$ | - |
| $g$ | vector of gains | $Q$ | - |
| $B$ | basis matrix | $Q \times P$ | - |
| $w_t$ | vector of node weights at time $t$ | $P$ | - |
| $W$ | matrix of node weights, across timesteps | $P \times T$ | - |

### A.4 Solving the linear inverse-problem (DENDRO Step 1)

#### A.4.1 Building the basis matrix *B*

The basis matrix *B* ∈ ℝ^*Q×P*^ is constructed so that each voxel is influenced by its *K* nearest nodes. Let 𝒦 denote the set of indices of the *K* + 1 nodes which are nearest to voxel *i* (the reason for using *K* + 1 nodes in this index set will become clear momentarily). Letting *x*_*i*_ ∈ ℝ^3^ denote the location of voxel *i*, and letting *g*_*j*_ ∈ ℝ^3^ denote the location of node *j*, we have

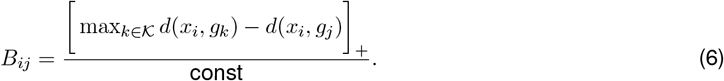

The distance *d*(·, ·) above is calculated using the path length of traversing nonzero voxels in the segmentation, and the constant in the denominator is chosen so that each row of *B* sums to 1. Scaling each row of *B* this way ensures that if voltage is constant across all nodes, then brightness will be constant across all voxels (because *B***1** = **1**). Our choice to use *K* + 1 nodes in the index set 𝒦 ensures that if node *j* is not in the set of the *K* nearest neighbors to voxel *i*, then *B*_*ij*_ will be zero. Thus each column of *B* forms a smoothly decaying basis function with local support. The hyperparameter *K* controls the size of each spatial basis function. We fixed *K* = 3 in all experiments.

#### A.4.2 Defining the microscope matrix *M*

The microscope matrix *M* ∈ ℝ^*D×Q*^ is defined using the equation for a gaussian beam. Letting *σ*(*z*) be the standard deviation of the (2D) gaussian psf at a distance *z* from the focal plane, we have:

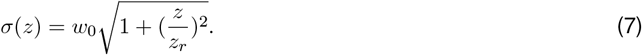

In practice, we build this matrix by looping over the non-background voxels in the segmented anatomy. For each voxel, we simply add a gaussian bump to the camera matrix, where the width of the bump is determined by the distance between the voxel and the focal plane according to Equation 7. We set the parameters *z*_*r*_ and *w*_0_ by matching them to the data by eye.

#### A.4.3 The DENDRO algorithm

The algorithm for solving the linear-inverse problem relies on simple alternating minimization to estimate the parameters of the model given in Equation 2. The full algorithm is given in Algorithm 1. Below, we discuss how each of the parameter updates are implemented.

##### Algorithm 1

DENDRO Alternating Optimization

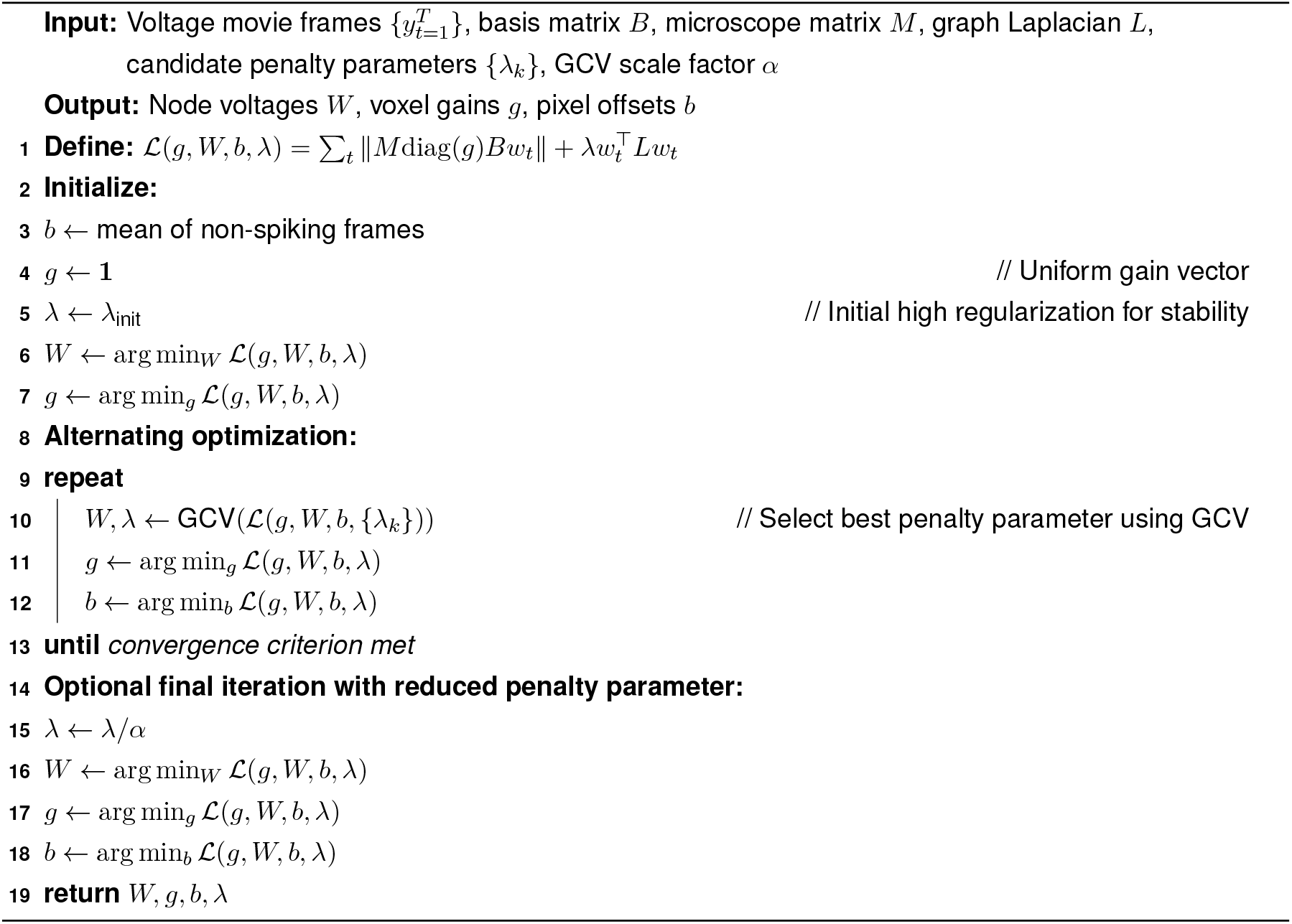

#### A.4.4 Generalized cross validation (GCV)

To lighten our notation, we write the DENDRO inverse problem for a single frame as 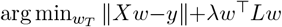. GCV offers an efficient approximation of the leave-one-out cross-validation error [Golub, Heath, and Wahba, 1979]. It requires computing the trace of the hat matrix *S* = (*X*^⊤^*X* + *λ*)^−1^*X*^⊤^*X*.

The matrix *S* is *D × D*, which in our cases would be prohibitive because voltages movies typically have 10k pixels. Using the cyclic property of the trace, we have

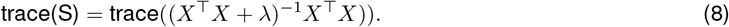

which only requires computing the diagonal entries of a *P × P* matrix.

#### A.4.5 Updating the gains *g* **and offsets** *b*

When solving for the gains, we impose nonnegativity and add a small ridge penalty:

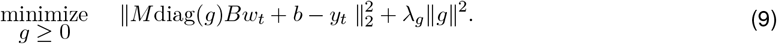

We solve the above problem using LBFGS-B and set *λ*_*g*_ = 0.001 for all real-data experiments. Updating the offsets *b* is done simply by setting it to the mean of the residual (though note that we initialize *b* using non-spiking frames). The simulated results do not include gains or offsets.

### A.5 Graph-based denoising architecture (DENDRO step 2)

Our architecture uses a combination of temporal and graph-based convolutions. The building block is a “graph-temporal blind-spot layer” (gtbs) which is shown in Figure 2e. Within each of these gtbs blocks are spatial blind-spot operations (Figure 2d,e, red squares) and temporal blind-spot operations (Figure 2d,e, blue squares). The temporal blind-spot blocks are implemented by a masked convolution, where the the center of the filter is held at zero.

The spatial blind-spot blocks are implemented as follows. Recall that *P* is the number of nodes in the graph. For a spatial blind-spot block with dilation *k* we first create a *k*-hop adjacency matrix *A*_*k*_(where two nodes are connected iff. they are *k* hops apart). Each spatial blind-spot block has a weight matrix 𝒲 of shape *C*_in_ *× C*_out_ (the number of input and output features respectively). Given a *P × C*_in_ input matrix *x*_*l*_, the spatial blind-spot operation forms the output *x*_*l*+1_ = *σ*(*Ax*_*l*_*W*) where *σ* is an activation function (Relu). The operation is broadcast along the time axis, so that in practice the input is a *P × C*_in_ *× T* tensor and the output is a *P × C*_out_ *× T* tensor.

We combine this block according to the architecture proposed by [Honzátko et al., 2020] (Figure S1). At the final layer of our network, we have a *P × C × T* (nodes, channels, timesteps) array. Until this point in the network, all operations are convolutional (i.e the network does not distinguish between nodes). We reasoned that it may be useful to break this symmetry, and so we use node-specific weights at the final layer. We define a final weight matrix with shape *P × C*. The output is then given by an element-wise multiplication with this weight matrix and sum along the channel dimension.

**Figure S1.**
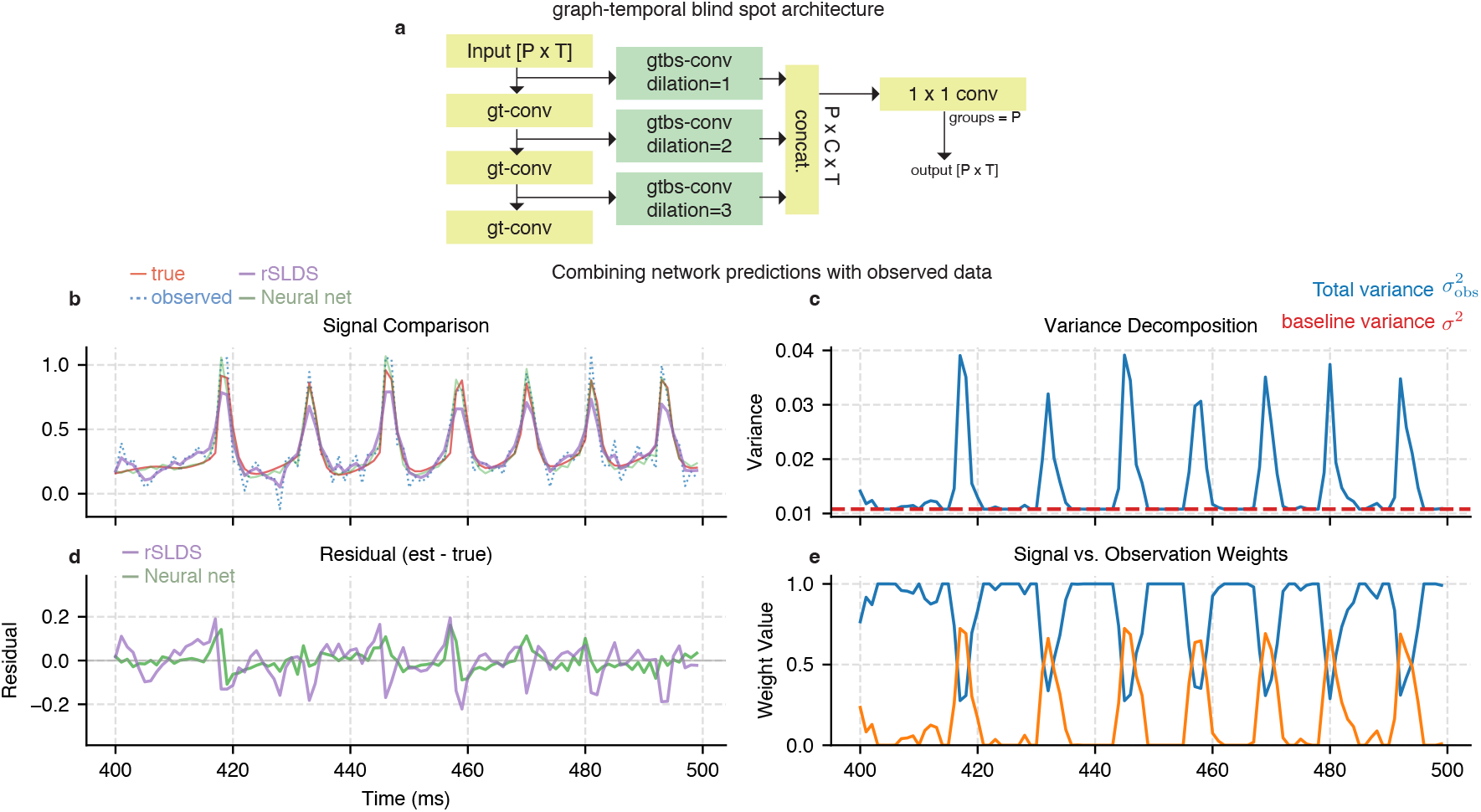
**a**, Graph-based blind-spot denoising architecture. Each block in green is a “graph-temporal blind-spot” layer. Each graph-temporal layer in yellow (“gt-conv”) follows the same structure as the gtbs layer, but without the blindspot. Each yellow block expands the receptive field, while the green blocks ensure the blind-spot property. **b**, Denoising example on single compartment data comparing rSLDS to a single-compartment neural net denoiser. **c**, Total variance predicted by the variance head of the network (blue trace) along with the baseline observation variance, computed by taking a percentile of the blue trace (in this case, the 10th percentile). **d**, Residuals (est -true) for rSLDS and neural net denoisers. **e**, Relative weighting of the network prediction (blue) and the observed data (orange). The observed data is upweighted during spikes.

#### A.5.1 Combining observations with network predictions

We found that the architecture above worked well overall, but still occasionally truncated spikes. We therefore implemented the gaussian smoothing model from [Laine et al., 2019] which adaptively mixes observed data and network predictions. Let the scalar 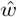 denote the observed noisy data at a given node and timestep (in practice, the output of DENDRO step 1), let *w* be the true denoised value, and let Ω_*w*_ be the surrounding context (nearby nodes at nearby timesteps). [Laine et al., 2019] propose a simple probabilistic model:

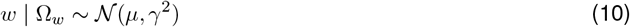

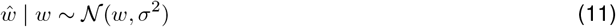

where *µ* represents the mean value after conditioning on the context, *γ*^2^ represents the variance in the true value after conditioning on context, and *σ*^2^ represents the observation noise. *µ* and *γ*^2^ vary for each node and timestep, whereas *σ*^2^ is a fixed observation noise variance (we allow different values of *σ*^2^ for each node).

Under this gaussian model, the total variance for an observed datapoint is the sum of the two variances, and the mean of an observed datapoint is equal to the mean when conditioning on context:

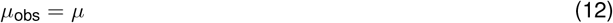

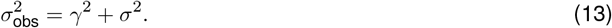

We train the network to predict *µ*_obs_ and 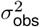, which vary across time and nodes. For each node, we extract its observation noise *σ*^2^ by taking a percentile of the total variance across time. We then form our final denoised estimates using a weighted combination of the network output and the observed data:

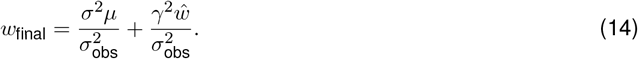

To illustrate how this process works in practice, we have provided an expanded single-compartment denoising example in Figure S1b-e. During spikes, the network outputs a heightened estimate of the total signal variance 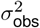 (Figure S1c). The network input and predictions are adaptively combined according to a time-varying convex combination. The increased total variance during spikes leads to a relatively higher weighting of the network input during spikes, reflecting that the spikes are more difficult to predict (Figure S1e).

Note that this is slightly different than the proposal from [Laine et al., 2019] – they instead train a separate network to predict *σ*^2^ and they train the variance head of their network to output *γ*^2^ rather than the total variance. We found that predicting the total variance resulted in more stable training, and further, it allows us to control the amount of smoothing that the network performs at test time, without re-training. Because we are setting *σ*^2^ as a percentile of the total variance output across time, increasing this percentile results in higher noise variance, and thus more smoothing (the network prediction is weighted more highly). Decreasing the percentile results in less smoothing (the input data is weighted more highly). For all simulated data experiments, we used the 95th percentile to calculate the noise variance. For real data, we used the 80th percentile.

#### A.5.2 Network training

All networks were trained with maximum likelihood as the objective, for 25 epochs, using the Adam optimizer with learning rate = 0.001, on a single NVIDIA GTX1080TI.

### A.6 Assessing performance on simulated data

#### A.6.1 GEVI model

To accurately capture the transformation from membrane voltage to observed fluorescence, we modeled the GEVI kinetics using a four-state model driven by the membrane potential *v*(*t*) (Figure 2a). We denote the states *G, M, N, R*. The four state model was chosen because it fit the data well rather than because of biophysical considerations.

The dynamics are governed by instantaneous transition rates between states. Transitions *G* ↔ *M, N* ↔ *R*, and *R* ↔ *G* are voltage-independent with rate constants *k*_*gm*_, *k*_*mg*_, *k*_*nr*_, *k*_*rn*_, *k*_*rg*_, *k*_*gr*_, while transitions *M* ↔ *N* are voltage-dependent. We parameterize the *M* → *N* and *N* → *M* transitions with sigmoidal rate functions

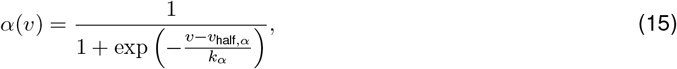

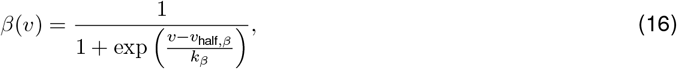

where *v*_half,*α*_, *v*_half,*β*_ are half-activation voltages and *k*_*α*_, *k*_*β*_ set the voltage sensitivity of the transitions. The resulting ODEs for the continuous-time state dynamics are

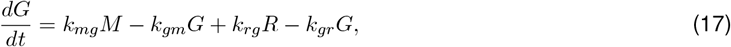

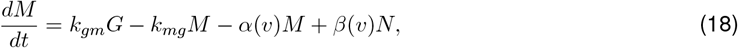

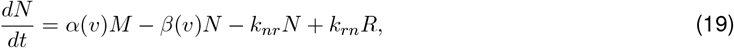

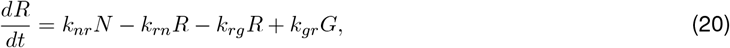

subject to the normalization constraint *G* + *M* + *N* + *R* = 1 at all times. We implemented this model as a current-free channel in Jaxley, so that it can be simulated as part of our single and multicompartment biophysical models.

Fluorescence is modeled as a linear readout of the state occupancies, plus a baseline term:

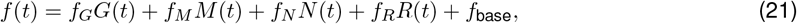

where *f*_*G*_, *f*_*M*_, *f*_*N*_, *f*_*R*_ are per-state gains and *f*_base_ captures background fluorescence. To account for the finite integration time of the camera, we introduce an additional “camera” fluorescence variable *f*_cam_(*t*), which low-pass filters the instantaneous indicator fluorescence:

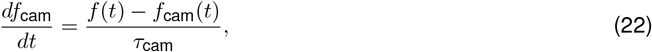

with time constant *τ*_cam_ set by the effective camera integration. During simulation, we record *f*_cam_(*t*) and treat it as the model’s prediction of the observed fluorescence trace.

#### A.6.2 GEVI Model fitting

The GEVI model has 16 trainable parameters, listed in Table 1. We fit these parameters using paired patch-clamp and fluorescence recordings (Figure 2a,b). Given a measured voltage trace *v*(*t*) and a simultaneously recorded fluorescence trace 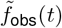, we simulate the indicator dynamics by driving the four-state model with *v*(*t*) and integrating the ODEs described above. During fitting, we resample the electrophysiology recordings to 40Khz and use this time step when integrating the GEVI model. This yields a predicted camera fluorescence trace *f*_cam_(*t*) along with the underlying state trajectories *G*(*t*), *M* (*t*), *N* (*t*), *R*(*t*). Because the imaging and electrophysiology traces are be sampled at different rates, we linearly interpolate *f*_cam_(*t*) to the fluorescence time stamps and define the loss as the mean-squared error between the predicted and observed fluorescence.

**Table 1.** Trainable parameters of the four-state GEVI model.

| Symbol | Meaning | Constrained $> 0$? |
| --- | --- | --- |
| $k_{gm}$ | $G \rightarrow M$ transition rate | Yes (Softplus) |
| $k_{mg}$ | $M \rightarrow G$ transition rate | Yes (Softplus) |
| $k_{nr}$ | $N \rightarrow R$ transition rate | Yes (Softplus) |
| $k_{rn}$ | $R \rightarrow N$ transition rate | Yes (Softplus) |
| $k_{rg}$ | $R \rightarrow G$ transition rate | Yes (Softplus) |
| $k_{gr}$ | $G \rightarrow R$ transition rate | Yes (Softplus) |
| $v_{\text{half},\alpha}$ | Half-activation voltage for $M \rightarrow N$ ("$\alpha$") transition | No |
| $v_{\text{half},\beta}$ | Half-activation voltage for $N \rightarrow M$ ("$\beta$") transition | No |
| $k_\alpha$ | Slope factor for $M \rightarrow N$ transition | No |
| $k_\beta$ | Slope factor for $N \rightarrow M$ transition | No |
| $f_G$ | Fluorescence gain for $G$ state | No |
| $f_M$ | Fluorescence gain for $M$ state | No |
| $f_N$ | Fluorescence gain for $N$ (bright) state | No |
| $f_R$ | Fluorescence gain for $R$ state | No |
| $f_{\text{base}}$ | Baseline fluorescence offset | No |
| $\tau_{\text{cam}}$ | Camera integration time constant | No |

We optimize *θ* using gradient-based optimization (L-BFGS) with automatic differentiation through the GEVI simulation via Jaxley [Deistler et al., 2024]. To enforce positivity of the transition rates, we reparameterize the rate constants via a softplus transform and optimize in the unconstrained space, while parameters such as *v*_half,*α*_, *v*_half,*β*_, *k*_*α*_, *k*_*β*_ and the fluorescence gains are left unconstrained. At the start of each fit, we initialize the state occupancies *G, M, N, R* at their steady-state values given the resting potential of the cell, by solving a linear system. After optimization converges, we fix the learned parameter set 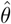 and use it for all subsequent simulations and validation experiments described in the main text.

#### A.6.3 Generating single-compartment data

All channel-based simulations were conducted using Jaxley [Deistler et al., 2024]. For the single-compartment simulations (Figure 2a,b) we used a Hodgkin-Huxley model, and added a simulated opsin channel (reversal potential at 0mV). We then drew samples from a gaussian-process (RBF kernel with lengthscale = 30ms) to excite our simulated opsin channel, which created spikes and smooth subthreshold waveforms. These simulated voltage values drove the GEVI model described above to produce simulated fluorescence traces.

#### A.6.4 Generating multicompartment data and simulated voltage videos

Multicompartment data was also generated using Jaxley, similar to the procedure described above. For maximum fidelity to our real-data experiments, we used segmentations (graphs and voxels) from real cells in our dataset (see subsection A.7 for details on segmentation). We then re-implemented the biophysical model from Park et al. [2025] in Jaxley. We drew a randomly fluctuating time-series from a GP, and used it to inject current into an apical dendritic compartment. As expected, our multicompartment model produced bAPs both with and without associated dendritic spikes, along with subthreshold dynamics. As in the single-compartment case, these simulated voltages then drove simulated fluorescence via the GEVI model. We treated these GEVI outputs as ground-truth node weights *w*_*t*,true_ (shown in Figure 3c, bottom).

To generate simulated voltage videos, we created a basis matrix *B* associated with the multicompartment model (1 basis function per compartment). This yields ground truth 3D voltage signals *v*_*t*,true_ = *Bw*_*t*,true_. These 3D voltage signals (really, 3D voxel fluorescence signals) can then be compared even when the underlying graph used to reconstruct the data departs from that used to generate the data (see next section).

We generated voltage videos by pushing our simulated 3D signals through a microscope model, then added multiplicative noise to each pixel:

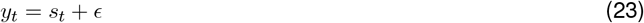

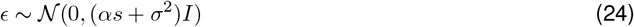

where *s*_*t*_ is the noiseless frame, *α* controls the amount of signal-dependent noise, and *σ*^2^ controls the signal-independent noise. We set the parameters *α, σ*^2^ to match the real data as follows. First, we observed that the full DENDRO pipeline offers excellent frame-level denoising, so we plotted total variance and noise variance for each pixel (Figure S3 left, middle). We then adjusted the noise parameters so that the noise variance of our simulated data (relative to the pixel means) matched that of the real data (Figure S3 right). This resulted in simulated videos which, by eye, were nearly indistinguishable in noise level from the real data (Supplemental video 1 shows a side-by-side comparison).

**Figure S2.**
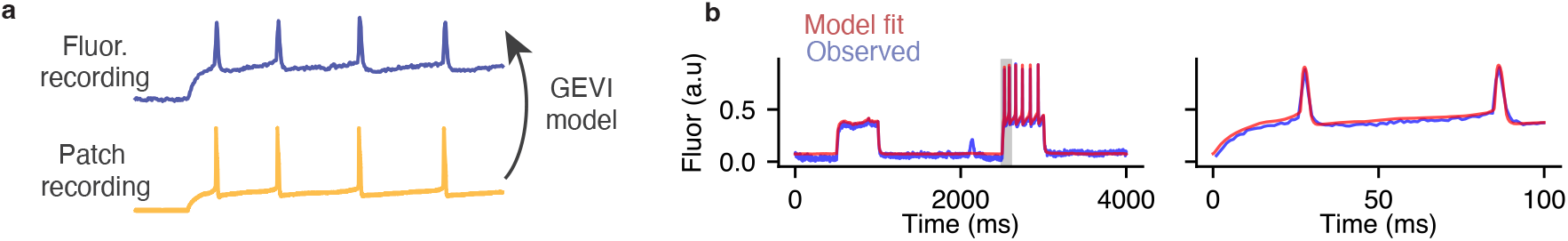
Four-state GEVI model captures the transformation from voltage to fluorescence. **a**, Four state GEVI model was trained on a paired patch-clamp plus fluorescence dataset. Parameters were learned to map observed voltage (from the patch clamp) to observed Voltron2 fluorescence. **b**, Left: observed vs. predicted fluorescence from the GEVI model during current steps. Patch recording is omitted for visual clarity. Right: zoom to the gray highlighted region in the left panel.

**Figure S3.**
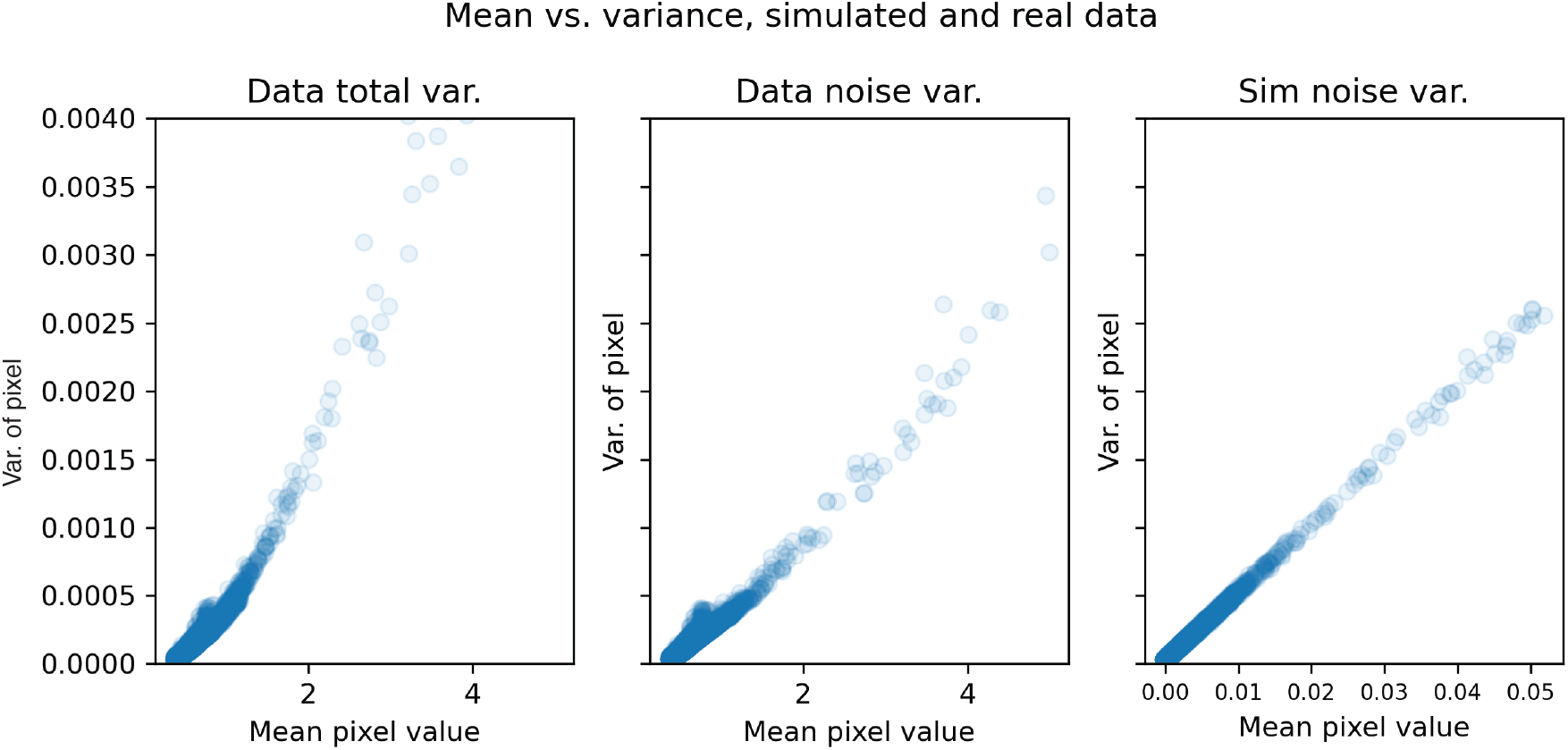
Calibrating simulated videos to match real data noise levels. Left: data total variance (each dot = one pixel) vs. mean pixel value. Middle: data noise variance vs. pixel mean. Right: Simulated noise variance vs. pixel mean. Note that the simulated noise variances span the same range as the real-data noise variances. The mean pixel values of the simulated data *do not* span the same range, because we not attempt to match the static fluorescence between simulated and real movies – i.e we ignore a scale factor between the real and sim data.

### A.7 Preprocessing acute slice datasets

Details of the experimental setup are given in [Park et al., 2025]. We segmented the 3D anatomy stacks using Simple Neurite Tracer [Longair, Baker, and Armstrong, 2011], which yielded a binary voxels array and an SWC file defining the skeletonized anatomy. We aligned the 3D anatomy (Z-projection) to the mean of the voltage movie by selecting control points on each image and finding an affine transform.

To preprocess the real data, we detrended each pixel using a b-spline (following the approach in [Buchanan et al., 2019]) to correct for photobleaching. We downsampled the voltage video (and aligned anatomy stack) by 4x in XY resulting in pixel sizes of 2.6*µ*m. For the simulated data results, we downsampled the anatomy stack by 2x in Z (yielding Z resolution of 2.0 *µ*m per voxel) and for the real data experiments we downsampled by 4x in Z (4 *µ*m per voxel).

### A.8 Extended real data results

To visualize our results on the real datasets, we generated similar comparison movies as we used for the simulated data (Supplemental videos 4 and 5). For each dataset, we provide: raw video, PMD video, DENDRO estimate, XY projection of inferred voltages, YZ projection of inferred voltages, residual (observed -estimated). We observed very little spatiotemporal structure in the residual, indicating that our model was not ignoring signal.

## Footnotes

1 We used the implementation of the rSLDS provided by SSM: https://github.com/lindermanlab/ssm

2 https://bit.ly/4rssK5h

