## Supplementary figures and images for "DENDRO: Recovery and denoising of whole-tree dendritic voltage from 2D voltage movies"

### supp_video4_realdata_cell1.tif

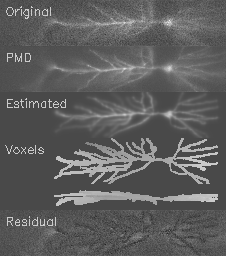

### supp_video5_realdata_cell2.tif

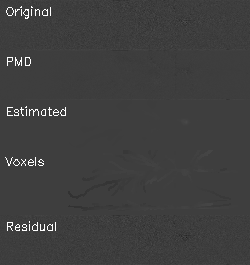
